# A novel texture backward masking method to locate critical recurrent processes in human vision

**DOI:** 10.64898/2026.06.17.732903

**Authors:** Netta Ollikka, Anni Bergström, Markku Kilpeläinen, Stéphane Deny

**Affiliations:** Department of Neuroscience and Biomedical Engineering, Aalto University, Finland; Department of Psychology, University of Helsinki, Finland; Department of Neuroscience and Biomedical Engineering, Department of Computer Science, Aalto University, Finland

## Abstract

Mounting evidence suggests that recurrent processes in the visual system play a critical role during challenging recognition tasks. Backward masking techniques have traditionally been used as a non-invasive method for studying recurrent processes: A mask follows the target image, presumably disrupting ongoing processes. However, these techniques have the limitation that they do not allow the identification of the *stage* of the visual system at which critical recurrent processes are taking place. Here, leveraging advances in texture synthesis *via* deep networks, and the approximate correspondence between stages of the visual system and layers of deep networks, we develop a novel psychophysics paradigm where masks with textures targeting different stages of the visual system follow the presentation of challenging images. In a series of experiments, we present objects to human subjects either for a short duration or in unusual poses, followed by a textured mask either designed to only target the early visual system, or the entire visual system. We find that both texture types equally affect recognition abilities, suggesting that recurrent processes in or towards early stages of the visual system are already recruited for these recognition tasks.

## 1 Introduction

Human vision is remarkably robust, enabling accurate object recognition under various conditions, such as unusual poses. Multiple lines of evidence converge to show that this robustness is supported by recurrent processes in the brain. First, the anatomy of the visual system and of the brain exhibit a highly recurrent structure (Felleman & Van Essen, 1991). Second, observational studies of brain activity in humans (e.g., Kietzmann et al., 2019) and non-human primates (e.g., Kar et al., 2019) provide evidence for recurrent processes within and/or across regions of the visual system. Third, computational studies using deep recurrent networks provide additional evidence that recurrence is critically involved in visual processing (e.g., Kietzmann et al., 2019; Seijdel et al., 2021). Finally, studies perturbing neural activity *via* transcranial magnetic stimulation (TMS) (e.g., Koivisto et al., 2011) have confirmed the importance of recurrent processes for challenging image recognition.

Backward masking is a widely used experimental technique for probing the temporal dynamics of visual processing. In this paradigm, a briefly presented target stimulus is followed shortly by a mask that can be, for example, a random checkerboard, a picture of an object, etc. While the mask arrives too late to interfere with the initial feedforward sweep, it is thought to interrupt the recurrent processing that typically follows. Perceptually, masking has little to no impact when participants view simple, non-ambiguous stimuli (Wyatte et al., 2012; Tang et al., 2018; Rajaei et al., 2019; Seijdel et al., 2021; Ollikka et al., 2025). However, it substantially impairs recognition of images that are degraded (Wyatte et al., 2012; 2014), occluded (Tang et al., 2018; Rajaei et al., 2019), rotated (Ollikka et al., 2025), or embedded in cluttered backgrounds (Seijdel et al., 2021); stimuli that are not self-explanatory and require additional processing, like recurrence. Neurophysiological studies have shown that early visual responses to the target remain intact under masking (Lamme et al., 2002; Fahrenfort et al., 2007; Seijdel et al., 2021), while later modulations, widely considered to reflect recurrence, are abolished (Lamme et al., 2002; Fahrenfort et al., 2007; Wyatte et al., 2012; Tang et al., 2018; Rajaei et al., 2019; Seijdel et al., 2021). Together, the neurophysiological and perceptual evidence suggests that backward masking perturbs or even interrupts recurrent processes which are critical to vision in challenging conditions. However, unlike the more invasive methods aforementioned, the backward masking paradigm is currently unable to locate *where* in the brain the critical recurrent processes are taking place. Most have manipulated the timing of target and mask (stimulus onset asynchrony, SOA) (e.g., Klein et al., 2026), while keeping the structure of the mask constant. As a result, they provide little insight into the anatomical origin of the disrupted recurrence. Others, who observed the effect of varying mask content, such as its intensity (e.g., Schiller, 1966) or pattern (e.g., Enns & Di Lollo, 1997; Hermens & Herzog, 2007), were not aiming at locating recurrent processes along the visual stream. Here, we propose a novel masking method to identify *where in the visual system* recurrence becomes critical for object recognition under challenging conditions.

The method proposed here relies on the established correspondence between convolutional neural networks (CNNs) for vision and the primate ventral stream. Both systems are organized hierarchically: early stages detect low-level features such as edges and contrast, while later stages extract more complex shapes and semantic information. Empirical work shows a large degree of functional correspondence between layers in CNNs and stages of the visual system, with early layers resembling activity in areas like V1 and V2, intermediate layers aligning with V4, and deeper layers matching responses in high-level areas such as the IT cortex, both spatially and temporally (Yamins & DiCarlo, 2016; Cichy et al., 2016a; b). This structural and functional similarity forms the theoretical foundation for using CNNs to target specific brain areas. Furthermore, several neurophysiological studies support the idea that different types of image structures engage distinct areas of the visual hierarchy (Freeman et al., 2013; Ziemba et al., 2016; Okazawa et al., 2017; Lieber et al., 2024). In particular, Freeman et al. (2013) examined responses to different textures in human and macaque visual cortex, and found that while V1 responses were similar across texture types, V2 already showed significantly greater responses to naturalistic textures. Okazawa et al. (2017) extended the analysis to V4, and found that the preference for naturalistic textures is even stronger in V4 than V2 (a preference also confirmed by Lieber et al. (2024) with photographs vs. scrambled photographs).

This study introduces a novel backward masking paradigm, in which the visual content of the mask is designed to target and disrupt recurrent processes at different levels of processing. We use structured noise masks generated using a CNN-based texture synthesis model (Gatys et al., 2015), designed to resemble either low-level visual features (e.g., edges and contrast) or high-level features (e.g., object-like structures). As argued in the previous paragraph, this approach should allow the masks to target different stages of the visual system. Indeed, a low-level mask should primarily activate V1 neurons, with activities decreasing along subsequent stages of the ventral stream, thus mainly interfering with recurrence at the early areas, such as intra-areal recurrence within V1 or inter-areal feedback projecting into V1 (Figure 1a.). In contrast, a high-level mask has the potential to disrupt recurrence at all levels, including intra-areal recurrence within higher areas (e.g., IT), or inter-areal feedback between mid- and high-level areas (e.g., V4–IT) (Figure 1b.). We applied our texture-based backward masking method to two types of challenging visual recognition tasks: scenes presented with a short exposure time (20 ms), and objects shown in unusual poses. In both these tasks, we found no effect of mask type on recognition accuracy or confidence. Two possible conclusions can be drawn from these results: (1) Early visual areas such as V1 are implicated in the critical recurrent process for recognizing these challenging images, or (2) Our low-level texture backmasks still residually perturb activity of high-level visual areas, making the localization of critical recurrent processes difficult with this method. We conclude this study with a critical analysis of these alternative explanations.

**Figure 1:**
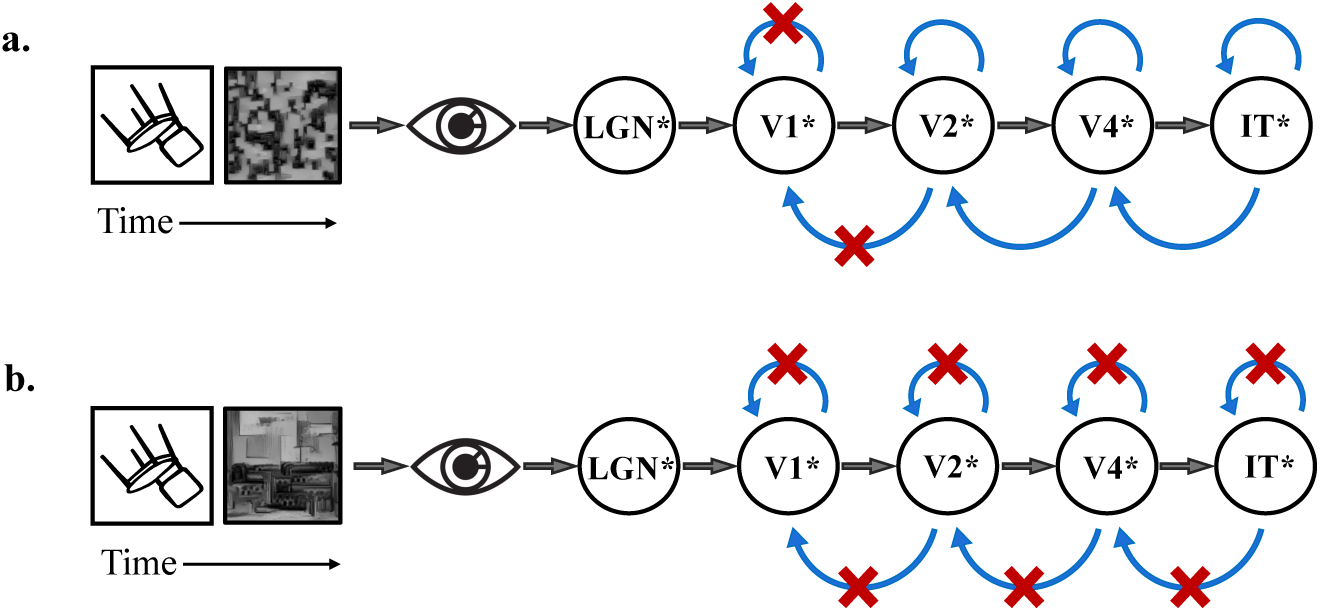
Illustration of the paradigm: Low- and high-level masks are hypothesized to disrupt recurrent processing at different levels of the visual hierarchy. Schematic illustration of the visual system (LGN, V1, V2, V4, IT) and recurrent interactions (blue arrows). Red crosses indicate the recurrent connections hypothesized to be disrupted by the mask. **a**. A target is followed by a *low-level* mask which primarily engages early visual cortex (e.g., V1) and is expected to interrupt recurrence locally within V1 and from higher areas projecting back to V1. **b**. A target is followed by a *high-level* mask which activates the entire visual hierarchy and is expected to disrupt recurrence more broadly, including intra-areal feedback in higher areas (e.g., IT) and inter-areal feedback between mid- and high-level areas (e.g., V4–IT). Asterisks (*) denote that the specific areas affected by respective masks are approximate.

## 2 Methods

We conducted a series of psychophysics experiments to investigate how visual masking affects object recognition under different sources of difficulty. Across experiments, we focused on two key challenges: limited viewing time and viewpoint variation. These challenges were selected because prior work suggests they require additional processing time and may engage recurrent computations (Mohsenzadeh et al., 2018; Ollikka et al., 2025). In addition, and in order to probe how different types of masks impact recognition, we systematically varied the structure of the mask (low-level vs. high-level) and its temporal dynamics.

We conducted five experiments, each involving a different set of participants. First, using synthetic objects, we presented target images in either upright or rotated poses for 60 ms, followed by a mask that was either low- or high-level (**Experiment 1**). In the rotated pose condition, which was found to be more challenging, we examined the differential effect of mask type (**Experiment 2**). Third, in the rotated pose condition, we also tested a condition where the target gradually transitioned into the mask (over 200 ms) rather than being followed by an abrupt onset (**Experiments 3 and 4**). Finally, using naturalistic images, we presented target images for either 60 ms or 20 ms followed by a mask that was either low- or high-level (**Experiment 5**). In the 20 ms condition, which was found to be more challenging, we examined the differential effect of mask type.

### 2.1 Observers

A total of 73 participants took part in the experiments. All participants had normal or corrected-to-normal vision and reported no history of neurological disorders (e.g., epilepsy). Participants came from diverse educational backgrounds, including psychology, engineering, and business, as well as participants without a background in higher education. Informed consent was obtained prior to participation. The study was conducted in accordance with the Declaration of Helsinki and was approved by the Research Ethics Committee of Aalto University (IRB: D/667/03.04/2022).

**Experiment 1** included 16 participants (8 female, 8 male; age range 22–57 years, mean age = 27.5 ± 8.2) and displayed conditions with upright poses and rotated poses, either followed by a mask or no mask. **Experiment 2** included 8 participants (all female; age range 20–45 years, mean age = 25.3 ± 8.0) and focused on rotated objects only, comparing low-level and high-level masks. **Experiment 3**, investigating the effect of gradual transitions between the target and the mask, included 8 participants (6 female, 2 male; age range 20–43 years, mean age = 29.3 ± 9.5). **Experiment 4**, a control condition for the third experiment was performed, where the target and mask were shown abruptly for 100 ms each. This experiment included 8 participants (3 female, 5 male; age range 20–58 years, mean age = 28 ± 11.5). **Experiment 5**, investigating recognition under extreme time limitations, included 33 participants (24 female, 9 male; age range 19–58 years, mean age = 30.9 ± 10.3).

### 2.2 Stimuli & Apparatus

#### 2.2.1 Target images

We created two stimulus sets for the object recognition tasks: one with synthetic 3D objects in unusual poses and one with naturalistic objects in canonical poses. The distribution of stimulus presentations across experiments is summarized in Table 1.

**Table 1:**
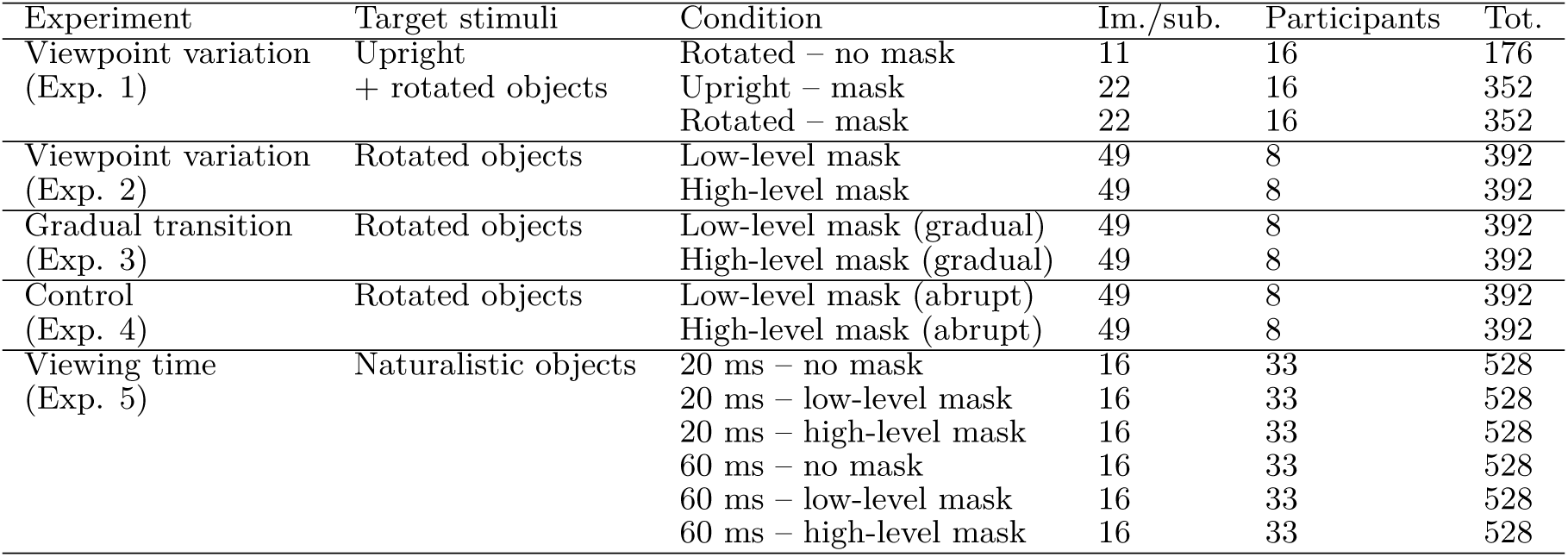
Distribution of stimulus presentations across experiments.

| Experiment | Target stimuli | Condition | Im./sub. | Participants | Tot. |
| --- | --- | --- | --- | --- | --- |
| Viewpoint variation<br>(Exp. 1) | Upright<br>+ rotated objects | Rotated – no mask | 11 | 16 | 176 |
|  |  | Upright – mask | 22 | 16 | 352 |
|  |  | Rotated – mask | 22 | 16 | 352 |
| Viewpoint variation<br>(Exp. 2) | Rotated objects | Low-level mask | 49 | 8 | 392 |
|  |  | High-level mask | 49 | 8 | 392 |
| Gradual transition<br>(Exp. 3) | Rotated objects | Low-level mask (gradual) | 49 | 8 | 392 |
|  |  | High-level mask (gradual) | 49 | 8 | 392 |
| Control<br>(Exp. 4) | Rotated objects | Low-level mask (abrupt) | 49 | 8 | 392 |
|  |  | High-level mask (abrupt) | 49 | 8 | 392 |
| Viewing time<br>(Exp. 5) | Naturalistic objects | 20 ms – no mask | 16 | 33 | 528 |
|  |  | 20 ms – low-level mask | 16 | 33 | 528 |
|  |  | 20 ms – high-level mask | 16 | 33 | 528 |
|  |  | 60 ms – no mask | 16 | 33 | 528 |
|  |  | 60 ms – low-level mask | 16 | 33 | 528 |
|  |  | 60 ms – high-level mask | 16 | 33 | 528 |

##### Synthetic 3D objects in unusual poses

were used to study the effects of viewpoint variation. This set builds on 51 objects introduced by Ollikka et al. (2025) and was expanded with 47 additional objects (98 total), following the same stimulus construction and selection procedure. The additional objects were obtained from Sketchfab, and rendered with OpenGL Alcorn et al. (2019); Abbas & Deny (2023). The objects were selected to belong to one of ImageNet categories. We discarded objects which were highly symmetrical (e.g., a ball), lacked meaningful rotated appearances (e.g., a restaurant), or were unlikely to be recognized by an average adult (e.g., a trimaran). For each object, images were generated in an upright pose with small viewpoint jitter (± 10^*°*^), as well as in 180 random out-of-plane poses sampled along the x-, y-, and z-axes. All objects were rendered on a uniform gray background and converted to grayscale to minimize contextual and color cues, so that recognition difficulty was primarily driven by viewpoint.

##### Naturalistic objects in canonical poses

consisted of 96 natural images obtained from the THINGS dataset (Hebart et al., 2019). This set was designed to probe recognition under extreme time-limited viewing while keeping the viewpoint canonical and preserving natural image structure (including background context).

In this condition, the temporal constraint accounted for the primary source of difficulty. These images were also converted to grayscale.

In order to get sensible alternative object categories for a two-choice object recognition task, we selected them with the help of a pretrained image classification model, Noisy Student EfficientNet (Xie et al., 2020), as described in detail in (Ollikka et al., 2025). Briefly, all created images of synthetic objects were given to the network to be classified. One upright image and one correctly classified rotated image were randomly selected per object. For the naturalistic images, all 96 images were classified. The correct alternative corresponded to the object’s ground-truth category. The incorrect alternative was defined as the network’s second-highest prediction. See examples of object images and response alternatives in Figure 2.

**Figure 2:**
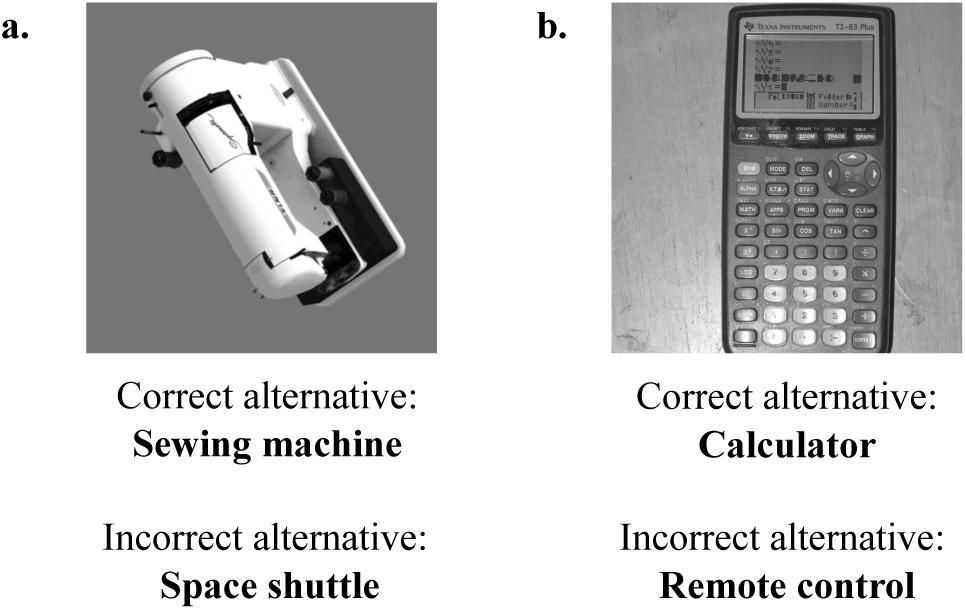
Example stimuli and answer alternatives used in the two-choice object recognition task. Each image is paired with two response options: the correct object alternative and the incorrect object alternative, which was derived from a deep neural network classifier. The incorrect alternative corresponds either to the model’s top prediction (if incorrect) or second-highest prediction (if correct). **a**. Example of a synthetic object in an unusual pose, used in the experiments investigating viewpoint variation. **b**. Example of a naturalistic object in a canonical pose, used in experiments investigating extreme time limitations.

#### 2.2.2 Mask images

We generated texture masks using the texture synthesis procedure of Gatys et al. (2015). This method uses a convolutional neural network (CNN), specifically VGG-19, and synthesizes images by matching feature statistics between a reference image and a generated texture. VGG-19 network consists of convolutional layers with ReLU nonlinearities and pooling. Each convolutional layer *l* contains *N*_*l*_ filters that produce *N*_*l*_ corresponding feature maps. Each feature map has *M*_*l*_ spatial locations. The activations of layer *l* can be arranged into a matrix 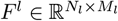, where 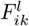 represents the activation of filter *i* at spatial location *k*.

The texture synthesis method is based on matching the Gram matrices of the reference image and the generated texture. The Gram matrix 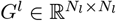 of either the reference image or generated texture for layer *l* is computed as

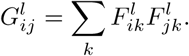

It measures correlations between feature maps across spatial locations. These correlations capture which features tend to activate together, independent of where these co-activations may occur. Matching the Gram matrices, therefore, preserves feature co-activation statistics while largely removing global shape information. Texture images are synthesized by initializing a random noise image and iteratively updating it so that its Gram matrices match those of a reference image at selected CNN layers. This optimization is performed using backpropagation, which adjusts pixel values to minimize the difference between the generated and reference image statistics. By choosing different layers of VGG-19 for this matching process, the synthesized textures can preserve either low-level features (such as edges and contrast) or more complex higher-order structures (Figure 3a.).

**Figure 3:**
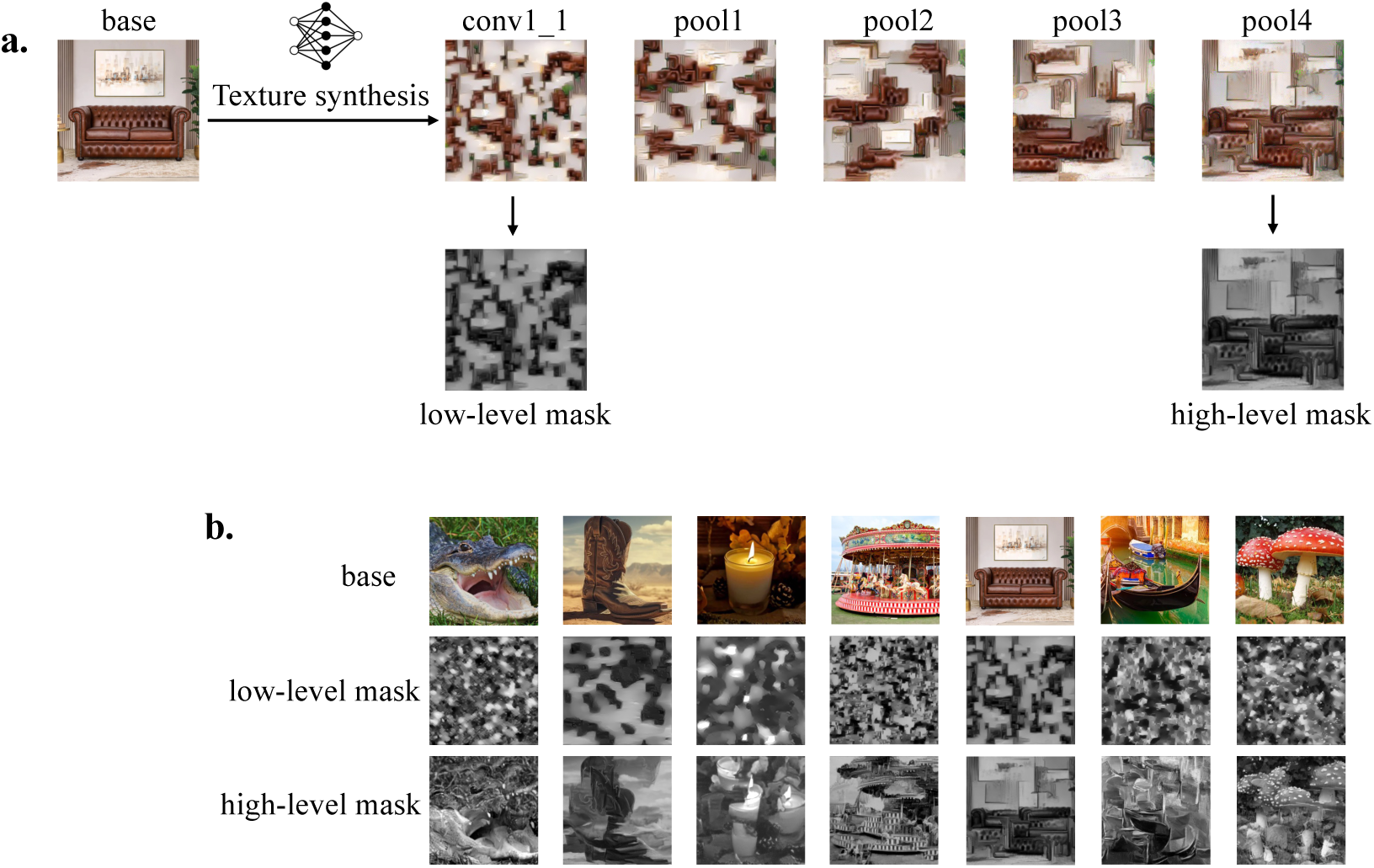
The texture masks were generated using CNN-based texture synthesis by Gatys et al.(2015). **a**. Masks were created by matching the Gram matrix statistics of selected VGG-19 layers, preserving feature correlations at a given level of the hierarchy while removing global shape information. Depending on the layers used, the masks capture either low-level features (e.g., edges and contrast) or higher-level structures. **b**. Examples of base images and the corresponding low- and high-level masks used in the experiments.

Following the texture synthesis method described above, we generated the texture masks used in the experiments (examples in Figure 3b.). The aim was to create masks containing either low-level or high-level visual feature structures. Low-level masks were generated by matching the Gram matrices computed from the earliest convolutional layer *conv1_1* using a 1 *×* 1 tiling configuration. In this configuration, feature statistics are matched across the entire image, thus these masks preserve local contrast and edge statistics. High-level masks were generated by matching the Gram matrices up to the deepest layer *pool4* using a 2 *×* 2 tiling configuration. Here, the image is divided into four spatial tiles, and texture statistics are applied within each quadrant as in Jagadeesh & Gardner (2022). This allows the masks to preserve some spatial information while still containing abstract and higher-order features. Masks were synthesized from several base images that were obtained from Google or THINGS database and that did not depict any of the objects used as targets in our experiments. See Appendix A for more details about mask creation.

#### 2.2.3 Apparatus

The experiments were implemented in MATLAB using Psychtoolbox (Kleiner et al., 2007). Visual stimuli were presented on a 22.5” VIEWPixx display with a resolution of 1920 × 1200 pixels, a 100 Hz refresh rate, and a viewing distance of 54 cm from the screen. The experiment room was dimly lit, and the head position was stabilized with a chinrest. The diameter of the object images and masks was 13.3 degrees of visual angle. Responses were collected using a standard keyboard. Response alternatives were in Finnish (native language of all participants).

### 2.3 Procedure

Before the experiments, participants completed a short training session to familiarize themselves with the task. The training trials used object images that were not included in the main experiments. All experiments followed the same general trial structure (Figure 4). Each trial began with a central fixation cross presented on a gray background for an unlimited duration. Participants initiated the trial when ready by pressing a button, after which a target image was briefly presented at the center of the screen. A texture mask was then presented either immediately after the target or with temporal overlap, depending on the condition (with a no-mask condition included in some experiments). Participants then performed a two-alternative object recognition task by selecting the best matching alternative from two written options, one of which was always correct. The position of the response options (left/right) was randomized across trials, and trial order was randomized for each participant. In **Experiments 2, 3 and 4**, participants additionally reported their confidence by indicating whether they were *sure* or *unsure* after the response (See confidence analysis in Appendix B). Each participant saw every target image only once.

**Figure 4:**
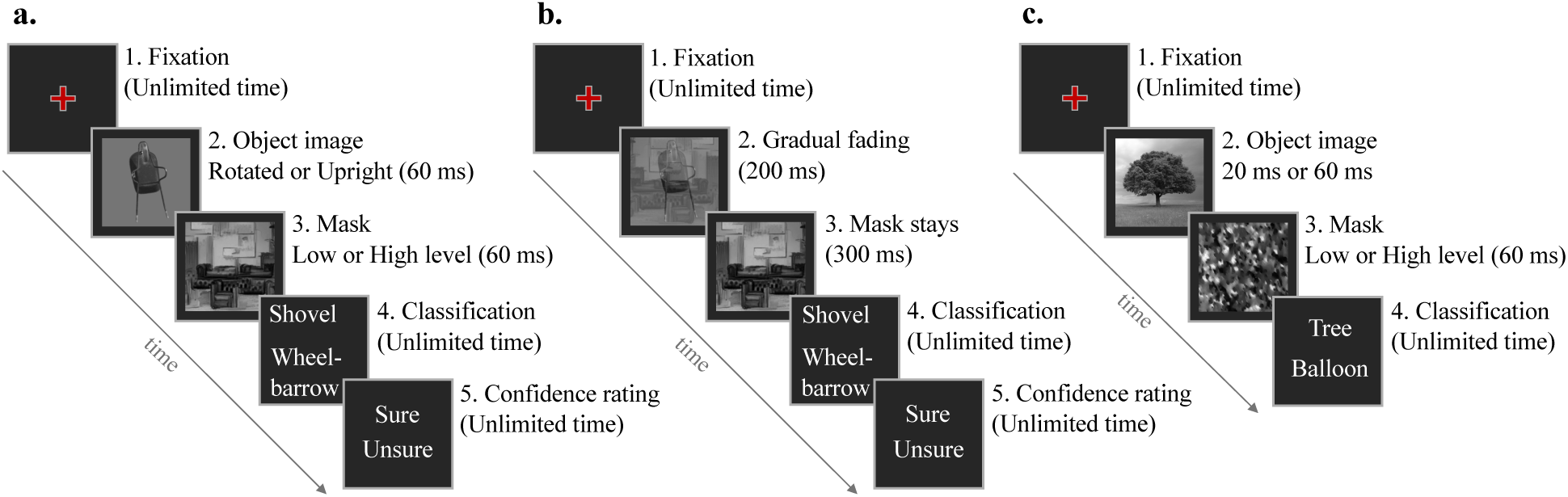
Experimental procedures used in this study. **a.**Viewpoint variation experiments (Exp. 1, 2 and 4). Each trial begins with a central fixation cross, followed by a target object (upright or rotated) presented for 60 ms. The target is then followed by either a low-level or high-level mask (60 ms). Participants perform a two-alternative object recognition task and provide a confidence rating; response time is unlimited. **b**. Gradual masking experiment (Exp. 3). The trial structure is similar, but the target gradually fades into the mask over 200 ms. The mask then remains for another 300 ms after the transition. **c**. Extreme time-limitation experiment (Exp. 5). After fixation, a naturalistic object image is presented briefly (20 ms or 60 ms), followed by a low-level or high-level mask (60 ms). Participants then perform a two-alternative object recognition task.

Stimulus timings differed across experiments. In **Experiment 1**, the target image, either upright or rotated, was presented for 60 ms followed by a 60 ms mask or no mask. In **Experiment 2**, the target image, always rotated, was presented for 60 ms followed by a 60 ms low- or high-level mask. In **Experiment 3**, the target image, always rotated, gradually transitioned into the mask (low- or high-level). In this condition, the transition began at stimulus onset and followed a sigmoid function over 200 ms, resulting in a continuous transformation from target to mask. After the transition was completed, the mask remained visible for an additional 300 ms. In **Experiment 4** (a control for Experiment 3) the target was presented abruptly for 100 ms followed by a 100 ms low- or high-level mask. This condition matched the temporal integral of stimulus intensity in the gradual condition. In **Experiment 5**, a naturalistic image was shown for either 20 ms or 60 ms, and followed by a 60 ms low- or high-level mask or no mask.

## 3 Results

We investigated how different texture masks affect human object recognition under challenging conditions that are thought to engage recurrent processing in the visual system. Participants performed an object recognition task in which briefly presented target images were followed by either a mask or no mask. Task difficulty was manipulated along two dimensions: viewing time (20 ms vs. 60 ms) and object viewpoint (upright vs. rotated). When present, masks consisted of either low-level or high-level texture patterns derived from a VGG-19 network, designed to disrupt recurrence in different stages of the visual hierarchy (see Methods). We first identify conditions that impair recognition performance and then examine whether mask type differentially affects performance under these conditions. Finally, we introduce a gradual fading paradigm, in which the target image transitions smoothly into the mask instead of being followed by an abrupt onset.

### 3.1 Identifying challenging conditions

First, we examined the effect of viewing time on object recognition (Figure 5). Participants viewed naturalistic images of objects in a canonical pose with a background for either 20 ms or 60 ms, followed by a 60 ms mask. Recognition accuracy differed substantially between the two exposure durations. For the 60 ms condition, mean accuracy was 83.7% ± 2.7% (95% CI), whereas for the 20 ms condition, accuracy dropped to 63.1% ± 3.9%, corresponding to a difference of approximately 20 percentage points (paired t-test, t(32) = 9.60, p = 6.1e-11). Next, we examined the effect of object rotation. Here, the participants viewed objects presented either in their upright orientation or in a rotated pose for 60 ms, followed by a 60 ms mask. Recognition performance was substantially higher for upright poses, with mean accuracy of 87.1% ± 2.9%, than for rotated objects, with mean accuracy of 67.6% ± 6.9%, corresponding to a difference of approximately 20 percentage points (paired t-test, t(15) = 4.85, p = 2.1e-4). Across experiments, the masks used were both low-level and high-level. However, to isolate the effect of task difficulty (time limitation and rotation), performance was averaged across mask types in the analyses reported above.

**Figure 5:**
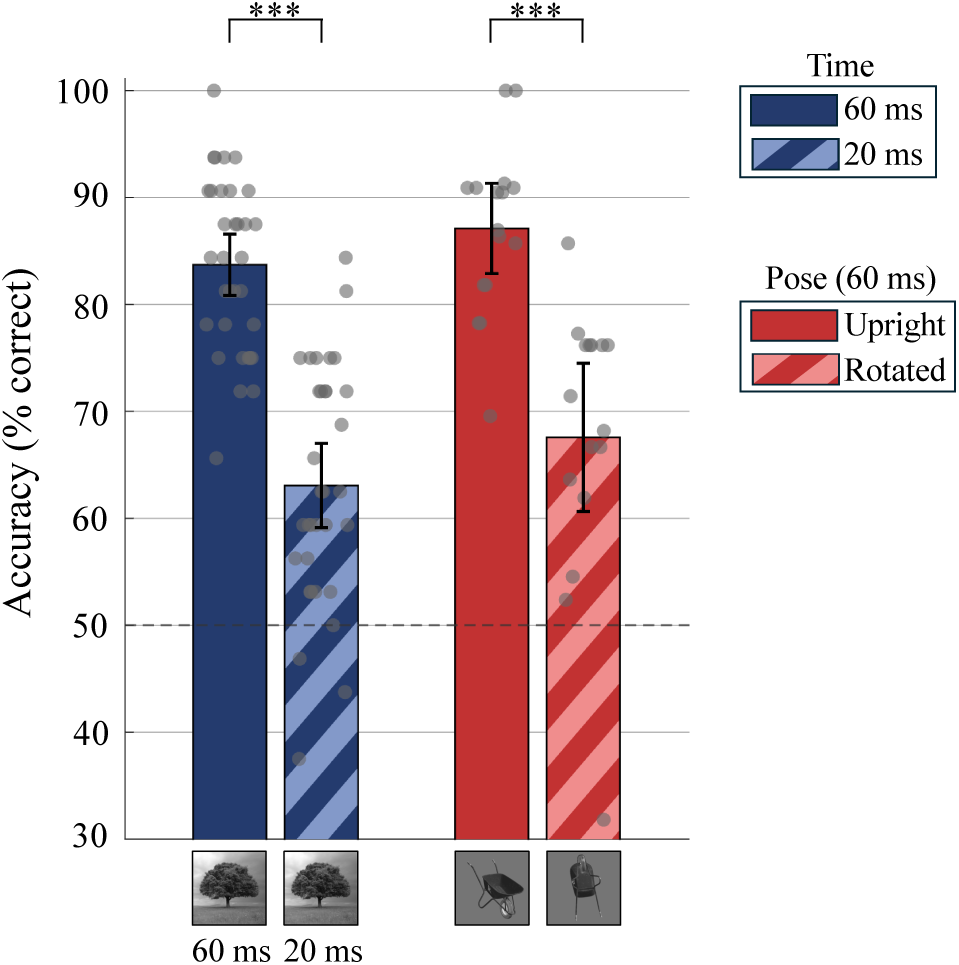
Identifying challenging conditions. *Blue bars:* Average performance in the extreme viewing time experiment (solid: 60 ms, striped: 20 ms). *Red bars:* Average performance in the viewpoint variation experi-ment (solid: upright, striped: rotated; both at 60 ms). Recognition accuracy decreased substantially under both manipulations. Gray dots represent individual participants, and error bars indicate 95% confidence intervals. Chance performance is 50%, and three stars (***) indicate highly significant differences (*p <* 0.001).

### 3.2 Comparable disruption from low- and high-level masks

We next examined whether mask type (no mask vs. low-level vs. high-level) differentially affected recognition performance under the two challenging conditions, extreme viewing time (20 ms) and object viewpoint (unusual rotation) (Figure 6). In the extreme viewing time experiment, in the no-mask condition, accuracy was nearly perfect, at 96.0% ± 1.3%. In the mask conditions, mean accuracies dropped to 64.0% ± 5.1% with low-level masks and to 62.1% ± 4.8% with high-level masks, and were significantly lower than in the no-mask condition (low-level: paired t-test, t(32) = 13.09, p = 2.12e-14; high-level: paired t-test, t(32) = 14.49, p = 1.32e-15). The difference between low-level and high-level mask accuracies was only 1.9 percentage points, which was not statistically significant. A similar pattern was observed in the unusual viewpoint experiment: when the rotated targets were not followed by a mask, recognition accuracy was high, at 90.3% ± 3.6%. When targets were followed by masks, accuracies dropped to 64.8% ± 11.5% for low-level masks and to 66.3% ± 5.8% for high-level masks, and were significantly lower than in the no-mask condition (low-level: unpaired t-test, t(22) = 6.16, p = 3.37e-6; high-level: unpaired t-test, t(22) = 8.11, p = 4.74e-8). The difference between low-level and high-level mask accuracies was only 1.5 percentage points, which was not statistically significant. Overall, recognition performance was comparably reduced across both mask types in both challenging conditions, indicating that low- and high-level masks produce similar levels of disruption.

**Figure 6:**
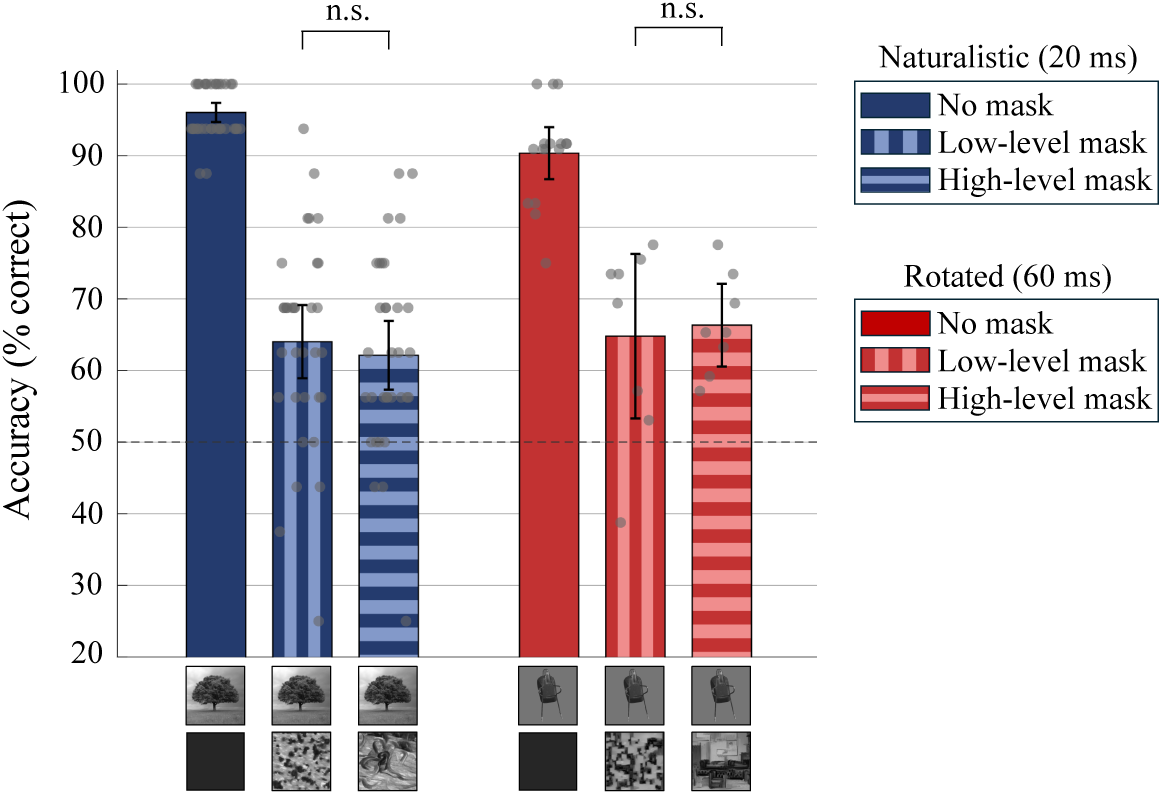
Comparable disruption from low- and high-level masks. *Blue bars:* Average performance in the extreme viewing time condition (naturalistic images, 20 ms). *Red bars:* Average performance in the viewpoint variation condition (rotated objects, 60 ms). Within each condition, bars show performance for no mask (left), low-level mask (middle), and high-level mask (right). Recognition performance was strongly impaired by masking in both conditions. However, low-level and high-level masks produced comparable levels of disruption, with no significant differences between them. Gray dots represent individual participants, and error bars indicate 95% confidence intervals. Chance performance is 50%, and “n.s.” indicates non-significant differences between mask types.

#### Interpretation

The comparable disruption observed for low- and high-level masks suggests that recurrence is not restricted to higher levels of the visual hierarchy. Low-level masks, which primarily contain edge- and contrast-level features, engage neurons in lower stages of the visual hierarchy but may not cause strong neural responses in higher-level areas like IT. In contrast, high-level masks may affect both early and later stages due to the hierarchical structure of visual feature processing. Thus, the similar disruptive effect of both mask types could imply the involvement of recurrence within the early visual cortex, such as V1.

### 3.3 From abrupt to gradual masking

One possible explanation for the similar effects of low- and high-level masks on recognition is the abrupt nature of the stimulus transitions in the standard backward masking paradigm. Abrupt onset and offset of the target and mask resemble a temporal step function, which likely elicits strong, broadband neural responses throughout the visual hierarchy. These global responses may override differences in the feature content of the masks and thus reduce their spatial specificity. To address this potential confound, we introduced a new masking technique in which the target image is gradually transformed into the mask. Specifically, the target was faded into the mask over 200 ms using a sigmoid function. This minimizes abrupt changes and ensures that the visual input changes continuously. As a control condition, we also included a conventional abrupt masking condition in which the target was presented for 100 ms and was immediately followed by a 100 ms mask. Importantly, both the gradual (200 ms sigmoid transition) and abrupt (100 ms target + 100 ms mask) conditions were matched in overall stimulus energy over time (i.e., the temporal integral of target image intensity was the same).

However, we again observed no significant differences between the low- and high-level mask conditions (Figure 7). Under the gradual condition, mean performance was 76.8% ± 9.1% for low-level masks and 75.0% ± 2.8% for high-level masks. These results were comparable to the abrupt condition, where mean performance was 76.0% ± 7.4% for low-level masks and 72.7% ± 7.6% for high-level masks.

**Figure 7:**
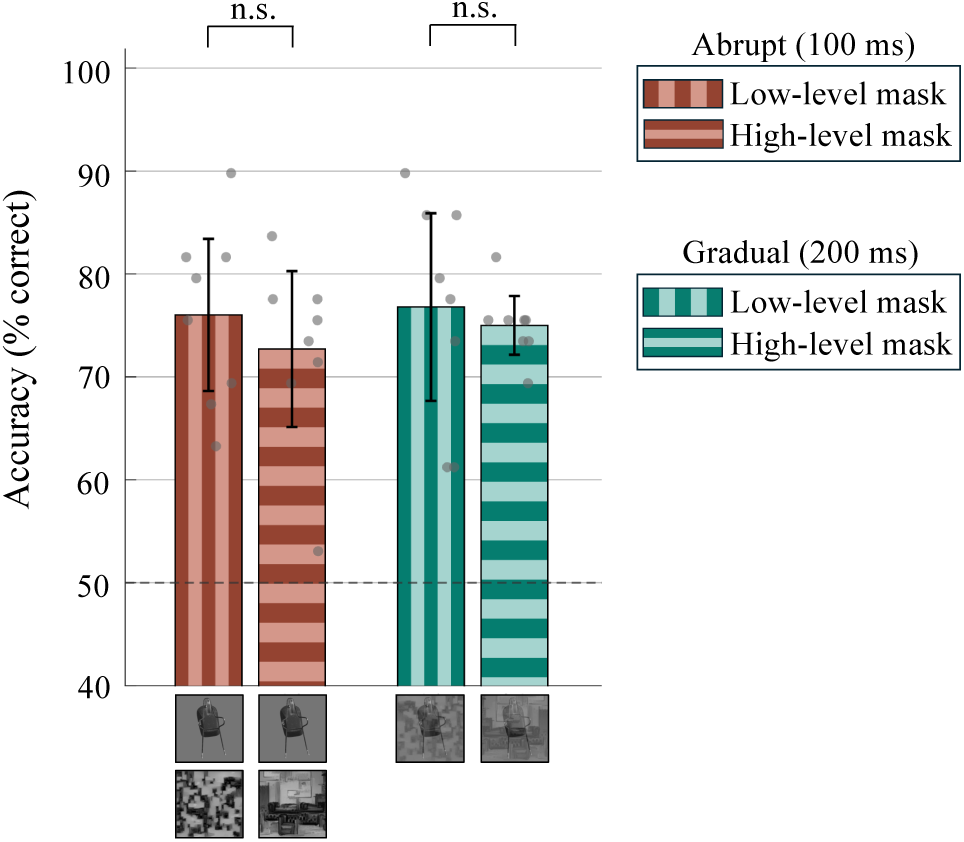
Comparable disruption under abrupt and gradual masking. *Orange bars:* Average performance in the abrupt masking condition (100 ms target followed by 100 ms mask). *Teal bars:* Average performance in the gradual masking condition (200 ms sigmoid transition from target to mask). Within each condition, bars show performance for low-level masks (vertical stripes) and high-level masks (horizontal stripes). Recognition performance did not differ between low-level and high-level masks in either condition. Gray dots represent individual participants, and error bars indicate 95% confidence intervals.

## 4 Discussion

We studied how different types of texture masks affect object recognition in a backward masking setting. In an attempt to perturb object processing at different levels of the human visual system, we used mask textures known to differentially activate different levels of convolutional neural networks (CNNs). Our results show that both low- and high-level masks impair object recognition, especially when the task is difficult (e.g., an uncommon pose or very short duration). We did not, however, find a significant difference between the mask types. This result could suggest that low-level visual areas are involved in critical recurrent processes for the recognition of challenging images (see Figure 1 for reasoning). This interpretation is in line with previous studies that have shown, using transcranial magnetic stimulation (TMS), the implication of V1 recurrence in diverse visual tasks (Koivisto et al., 2011; Roelfsema & de Lange, 2016; Hurme et al., 2019; Li et al., 2019). We address potential caveats to this conclusion in the following discussion.

### 4.1 Backward masking with different mask types

Studies using the backward masking paradigm have identified various functions of mask delay and mask type (reviewed by Breitmeyer & Ogmen (2000)). In the vast majority of cases, the mask has a substantial effect if presented immediately after the target image, as in the present study. Particularly relevant for the current study is the experiment by Bachmann et al. (2005). They used face images as targets and face images or noise as masks. Further, they varied the spatial resolution of the masks. They found that when mask spatial resolution was very low (making face and noise masks very similar), masking strength did not differ between the mask types. Differences (although not dramatic) in masking magnitude appeared with increasing spatial resolution of the mask. This pattern of results is probably at least partially due to cognitive mechanisms. For example, the target could be partially confused with the mask (which the authors themselves also suggest). We tried to avoid such a cognitive confusion. Our targets were never intact objects and never based on objects in a same category as the target. Although we did not test this directly, we believe that even our high-level masks were rarely recognizable to the observers, especially at such a short presentation time.

Our approach, in turn, comes with the caveat that while one can be certain that a fully recognizable face image and white noise differentially activate different levels of the visual system (Grill-Spector et al., 2017), we cannot be equally certain that our specific mask types do so. This caveat could explain why we observed no masking strength differences between the mask types. Our mask types, after all, were designed to differentially activate different layers of CNNs. We do not believe this approach is categorically misguided, as evidence from both humans (Cichy et al., 2016a; b) and non-human primates (Yamins et al., 2014; Yamins & DiCarlo, 2016) suggests considerable hierarchical similarity through the processing stages. More specifically, however, our set of mask levels—one intended to strongly activate a low processing level (e.g., V1) and another intended to additionally activate a level higher up in the visual stream (e.g., IT)— may be too sparse. We may be missing a mask that differentially activates a critical processing step. In addition, despite the overall hierarchical similarity, there may be substantial human-CNN differences in the algorithmic solutions within areas (Klein et al., 2026), which may make a specific mask type ineffective in selectively perturbing specific visual cortices (although see evidence to that regard in Introduction).

If we had observed stronger masking with the high-level masks, it would suggest that the important recurrent processes occur after V1. This follows the logic that masks should be the most effective when they strongly activate the neural level where the recurrence necessary for object recognition occurs. Just as V1 neurons practically ignore uniform luminance, higher-level neurons prefer complex textures over simple noise (Freeman et al., 2013; Okazawa et al., 2017). Therefore, if critical recurrence mainly occurred at a high processing stage, high-level texture masks should have caused greater perturbation. The equal effectiveness of the mask types instead suggests that at least some critical recurrent processes already occur at a lower level of the hierarchy (e.g., V1).

### 4.2 Masking is much stronger with challenging images

The current experiments clearly demonstrated that challenging targets (rotated or of short duration) are more strongly masked than easier targets. This finding adds to earlier studies (Bruchmann et al., 2010; Tang et al., 2018; Seijdel et al., 2021; Ollikka et al., 2025; Klein et al., 2026) on what makes the target stimulus particularly susceptible to masking. Many of the earlier findings can be explained by lower signal to noise ratio of the target leading to stronger masking (see Tsukamoto et al. (1990)). Our finding that objects in non-canonical poses are more susceptible to masking, cannot. Instead, it could be that objects in a canonical pose can be sufficiently processed in a purely (or mostly) feedforward manner, whereas more rotated objects require recurrent processing, which the subsequently arriving mask signal extinguishes. This would be in line with earlier results suggesting that object recognition with challenging images relies especially on recurrent processing (Rajaei et al., 2019; Kar et al., 2019).

### 4.3 Perspective

The level where recurrence processes critical for object recognition occurs has so far been mainly measured with TMS in humans and electrophysiology in animal models. Our study potentially provides a new, non-invasive and economical paradigm to locate critical recurrent processes, by-passing the need for more demanding and expensive experiments.

## Acknowledgments

We thank Noora Viljanen for additional data collection, and Felix Wichmann and Thomas Klein for an idea of experiment (gradual fading). This work has been supported by a Finnish Cutural Foundation (FCF) grant to N.O., and a Research Council of Finland (RCF) grant to S.D. under the Project: 3357590. Parts of this work were performed as part of the Master’s thesis of N.O. (Ollikka, 2025).

## Appendix

## A Texture masks

Masks were synthesized from several base images that varied across the experiments. In **Experiment 1**, we used 10 base images: *alligator, boot, candle, carousel, conch, couch, fountain, goldfish, gondola, and mushroom*. We created both low- and high-level masks with 1×1 tiling configuration. In **Experiments 2, 3 and 4**, we used 7 base images: *alligator, boot, candle, carousel, couch, gondola, and mushroom*. Here, the low-level masks had 1×1 tiling and high-level masks 2×2 tiling. In **Experiment 5**, we used 10 base images: *cow, flamingo, motorcycle, owl, snake, sunflower, typewriter, watering can, wolf, and zebra*. Here, the low-level masks had 1×1 tiling and high-level masks 2×2 tiling. The base images were always different from the objects used as experimental targets. For each base image, we generated multiple low-level and high-level masks and converted all masks to grayscale to match the object stimuli. In **Experiments 2, 3 and 4**, masks were normalized to mean luminance and root-mean-square (RMS) contrast. Because of the progressive refinement of our experimental design, this normalization was not applied to masks used in **Experiments 1 and 5**. We did not see any differential effect between normalized and non-normalized masks, making our conclusions robust to this design choice.

## B Signal detection theory (SDT) for confidence

Signal detection theory (SDT) provides a framework for separating sensitivity (the ability to detect a signal) from response bias (the tendency to favor one response over the other) (Stanislaw & Todorov, 1999; Macmillan, 2014). Here, SDT was applied to binary confidence judgments (confident vs. not confident) to assess whether the individual was able to discriminate between their correct and incorrect decisions. For this analysis, correct trials were treated as signal (*S*_2_) and incorrect trials as noise (*S*_1_), with a confident response indicating that the signal was present.

We computed the standard SDT rates: the hit rate, *H* = *P* (confident | correct) and the false-alarm rate, *F* = *P* (confident | incorrect).

Metacognitive sensitivity was quantified by

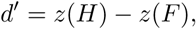

where *z*(·) = Φ^*−*1^(·) is the inverse of the standard normal cumulative distribution function (CDF). A larger *d*′ indicates that confidence is reported more often on correct than incorrect trials. A value of *d*′ = 0 indicates chance-level discrimination, i.e., the participant reports confidence equally often in both cases.

Confidence bias was quantified by

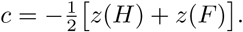

Values of *c <* 0 indicate a liberal tendency to report confidence, i.e., participants are overconfident. Values of *c >* 0 indicate a conservative tendency, i.e., participants are underconfident. A value of *c* = 0 indicates no bias.

We did not observe any significant differences between low- and high-level masks in either metacognitive sensitivity (*d*′) or confidence bias (*c*) (Figure 8). Participants were generally able to report higher confidence for correct than incorrect responses across conditions (positive *d*′). However, no significant differences in *d*′ were found between any condition (e.g., abrupt 60 ms vs. gradual 200 ms). In contrast, some differences were observed in confidence bias (*c*) across viewing conditions. Specifically, bias differed between the abrupt 100 ms and abrupt 60 ms conditions in both the low (0.0676 vs. 0.8827, *p* = 0.0036) and high (0.1831 vs. 0.8589, *p* = 0.0175) mask conditions. There was also a significant difference between abrupt 100 ms low (0.0676) and gradual 200 ms low (0.5476) with a p-value of 0.045. These results suggest that in the abrupt 100 ms condition, participants exhibited a weaker bias toward answering “not confident” compared to other conditions.

**Figure 8:**
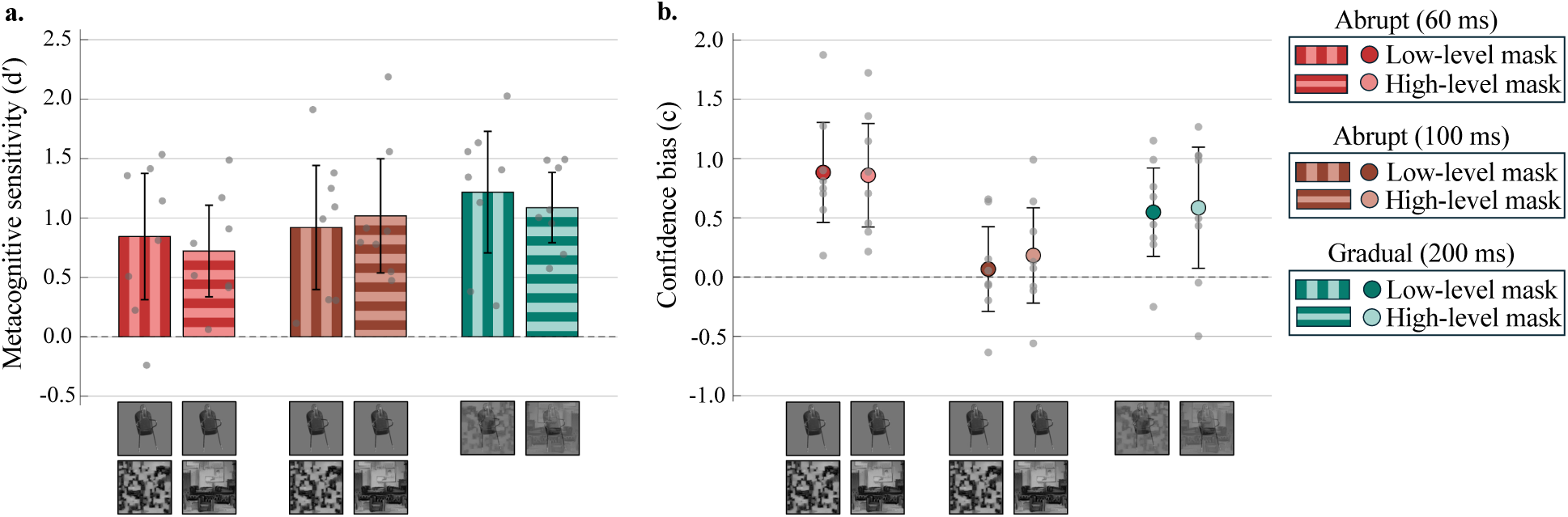
Metacognitive sensitivity and confidence bias across masking conditions. **a.**Metacognitive sensitivity (*d*′) across abrupt 60 ms (red), abrupt 100 ms (orange), and gradual 200 ms (teal) masking conditions. Vertical bars indicate low-level masks and horizontal bars indicate high-level masks. **b**. Confidence bias (*c*) across the same conditions. Darker circles indicate low-level masks and lighter circles indicate high-level masks. No significant differences were observed between low-level and high-level masks in either metacognitive sensitivity or confidence bias. Gray dots represent individual participants, and error bars indicate 95% confidence intervals. Dashed lines indicate the unbiased reference values (*d*′ = 0 and *c* = 0).

